# Regulation of dopamine-dependent transcription and cocaine action by *Gadd45b*

**DOI:** 10.1101/2020.05.01.072926

**Authors:** Morgan E. Zipperly, Faraz A. Sultan, Guan-En Graham, Andrew C. Brane, Natalie A. Simpkins, Lara Ianov, Jeremy J. Day

**Author notes:** Correspondence to Jeremy Day ( |).

## Abstract

Exposure to drugs of abuse produces robust transcriptional and epigenetic reorganization within brain reward circuits that outlives the direct effects of the drug and may contribute to addiction. DNA methylation is a covalent epigenetic modification that is altered following stimulant exposure and is critical for behavioral and physiological adaptations to drugs of abuse. Although activity-related loss of DNA methylation requires the *Gadd45* (Growth arrest and DNA-damage-inducible) gene family, very little is known about how this family regulates the activity of brain reward circuits or behavioral responses to drugs of abuse. Here, we combined genome-wide transcriptional profiling, pharmacological manipulations, electrophysiological measurements, and CRISPR tools with traditional knockout and behavioral approaches in rodent model systems to dissect the role of *Gadd45b* in dopamine-dependent epigenetic regulation and cocaine reward. We show that acute cocaine administration induces rapid upregulation of *Gadd45b* mRNA in the rat nucleus accumbens, and that knockout or site-specific CRISPR/Cas9 gene knockdown of *Gadd45b* blocks cocaine conditioned place preference. *In vitro*, dopamine treatment in primary striatal neurons increases *Gadd45b* mRNA expression through a dopamine receptor type 1 (DRD1)-dependent mechanism. Moreover, shRNA-induced *Gadd45b* knockdown decreases expression of genes involved in psychostimulant addiction, blocks induction of immediate early genes by DRD1 stimulation, and prevents DRD1-mediated changes in DNA methylation. Finally, we demonstrate that *Gadd45b* knockdown decreases striatal neuron action potential burst duration *in vitro*, without altering other electrophysiological characteristics. These results suggest that striatal *Gadd45b* functions as a dopamine-induced gene that is necessary for cocaine reward memory and DRD1-mediated transcriptional activity.

---

ADDICTION IS AN increasingly prevalent problem in the United States, associated with progressively higher rates of morbidity and mortality. Experience with drugs of abuse results in significant transcriptional and epigenetic alterations in brain reward circuits that support both synaptic and behavioral plasticity, outlasting the direct effects of the drug and contributing to the development of addiction [1–4]. Despite their various mechanisms of action, one common feature of all drugs of abuse is that they act upon the mesocorticolimbic dopamine (DA) system, which includes the ventral tegmental area (VTA), the nucleus accumbens (NAc), and the prefrontal cortex [2,5]. Addictive substances exert their rewarding effects in part by elevating DA concentrations in the NAc, a brain region integral to reward learning, as well as the development and maintenance of addiction [6–8]. This increase in DA modifies neuronal function by usurping mechanisms underlying adaptive forms of learning, activating signaling cascades and transcriptional programs that contribute to the reinforcing properties of drugs of abuse and leading to drug-evoked synaptic plasticity [2,6,8–12].

Epigenetic mechanisms include post-translational modification of histone proteins and methylation of cytosine-phospho-guanine (CpG) dinucleotides within DNA, which alter chromatin structure and transcription factor activity to modulate gene expression [1,4,13]. DNA methylation is catalyzed by DNA methyltransferases (DNMTs), and promoter hypermethylation is traditionally associated with transcriptional silencing [4]. Previous work has demonstrated upregulation of DNMT3A and DNMT3B (proteins which are required for *de novo* methylation) in the NAc 4 hours following acute cocaine treatment; by 24 hours, these levels decreased below baseline [4,14,15]. However, a more recent study examining methylation machinery in the NAc found decreased DNMT3A 30 minutes following acute cocaine treatment and increased levels of the protein at 24 hours [16], suggesting complex temporal dynamics in regulation of DNA methylation machinery. Intriguingly, both DNMT inhibition and chronic methyl supplementation blocked the expression of cocaine-induced locomotor sensitization, despite their opposing effects on DNA methylation [4,14]. Furthermore, intra-cranial delivery of the DNA methylation inhibitor RG108 resulted in enhanced cocaine conditioned place preference (CPP) when targeted to the NAc [15], but impaired associative reward learning when delivered to the VTA [17]. Following cocaine reward, differential hypo- and hyper-methylation events have been observed at specific gene loci [4,14,17]. Together, these findings suggest that DNA methylation dynamics are critically important for drug- and reward-related behaviors, and highlight the importance of identifying the underlying molecular mechanisms that contribute to these changes.

While numerous studies have demonstrated that DNA methylation marks are relatively stable and mediate long-term transcriptional regulation, the specific mechanisms that regulate CpG demethylation remain unclear. Previous studies have characterized the role of ten-eleven translocation (TET) enzymes in the conversion of 5-methylcytosine (5mC) to 5-hydroxymethylcytosine (5hmC) and additional oxidation that results in base excision and replacement [18–20]. In the brain, *Gadd45b*, a member of the Growth arrest and DNA-damage-inducible gene family, is associated with decreased CpG methylation in gene regulatory regions and is a critical for activity-dependent DNA demethylation, potentially working in concert with TET and thymine-dependent glycosylase proteins [18,21–24]. Moreover, *Gadd45b* is upregulated in response to neuronal activity in multiple brain regions and plays a key role in experience-dependent learning [21,22,25,26]. These results demonstrate that *Gadd45b* may link experience-dependent neuronal activity and downstream regulation of DNA methylation states.

However, despite enrichment of *Gadd45b* in the striatum [25–27] and clear links between drug-related transcriptional changes and DNA methylation, the role of *Gadd45b* in this process has not been examined. Here, we report that *Gadd45b* functions as an immediate early gene (IEG) in the rat NAc following acute cocaine reward or exposure to cocaine-paired contexts *in vivo*. Furthermore, *Gadd45b* disruption attenuates cocaine-paired place preference, suggesting that *Gadd45b* action is required for cocaine reward memory. Using a primary rat striatal neuron culture system, we demonstrate that DA-induced increases in *Gadd45b* require DRD1, Mitogen-activated protein kinase kinase (MEK), and cAMP response element binding protein (CREB) signaling. Furthermore, knockdown of *Gadd45b in vitro* results in the downregulation of IEGs and of genes implicated in dopaminergic synapse function and drug addiction and also abolishes changes in DNA methylation status following DRD1 activation. Intriguingly, while we observe significant transcriptional and behavioral alterations following *Gadd45b* manipulation, electrophysiological responses to DRD1 stimulation remain unchanged. Together, these results characterize *Gadd45b* as a DA-induced IEG in the striatum that is critical for transcriptional and behavioral adaptations to cocaine.

## Materials and Methods

### Animals

All experiments were performed in accordance with the University of Alabama at Birmingham Institutional Animal Care and Use Committee. Sprague-Dawley timed pregnant dams and adult male rats (90-120 days old) were purchased from Charles River Laboratories (Wilmington, MA). Dams were individually housed until embryonic day 18 (E18) for cell culture harvest. Male adult rats were co-housed in pairs in plastic filtered cages with nesting enrichment in an AAALAC-approved animal care facility maintained between 23-24°C on a 12hr light/dark cycle with ad libitum food (Lab Diet Irradiated rat chow) and water. Bedding and enrichment were changed weekly by animal resources program staff. Animals were randomly assigned to experimental groups. *Gadd45b* knockout mice for the cocaine conditioned place preference experiment were bred at the University of Alabama at Birmingham on a B6:129VJ background, as described previously [27,28]. Male wild-type and mutant offspring from heterozygotes were used for behavioral testing. Mice aged 2-6 months were individually housed at least 3 days prior to the start of behavioral testing. Heterozygotes were bred with C57BL/6 wild-types for at least six generations to generate back-crossed mice. All animals were handled for 3-5 days prior to behavioral testing.

### Drugs

Cocaine hydrochloride (C5776, Sigma-Aldrich, St. Louis, MO) was dissolved in sterile 0.9% sodium chloride and injected intraperitoneally (i.p.) at a dose of 10 mg/kg or 20 mg/kg for cocaine locomotor sensitization and conditioned place preference testing. Cocaine solution was made fresh immediately before behavioral testing and was protected from light.

For *in vitro* experiments, drugs were diluted in Neurobasal medium (Invitrogen, Waltham, MA) in a sterile hood immediately prior to treating cultured neurons. Dopamine hydrochloride (1µM; H8502, Sigma-Aldrich) was dissolved in Neurobasal medium. R(+)-SCH-23390 hydrochloride (D054, Sigma-Aldrich) and R(+)-SKF-38393 hydrochloride (S101, Sigma-Aldrich) were dissolved in sterile Milli-Q water and diluted to 1µM in Neurobasal medium. (-)-Quinpirole hydrochloride (Q102, Sigma-Aldrich) was dissolved in sterile Milli-Q water and diluted to 1µM in Neurobasal medium. Forskolin, 7-Deacetyl-7-[O-(N-methylpiperazino)-gamma-butyryl]-dihydrochloride (20µM; 344273, EMD Millipore, Billerica, MA) was dissolved in sterile Milli-Q water, then diluted in Neurobasal medium. For MEK inhibitor experiments, U0124 (Millipore, 662006) and U0126 (662005, Millipore) were dissolved in DMSO (D12345, Invitrogen) and diluted with Neurobasal medium to 1µM. The CREB inhibitor 666-15 (5661, Tocris, Minneapolis, MN), also called 3-(3-Aminopropoxy)-N-[2-[[3-[[(4-chloro-2-hydroxyphenyl)amino]carbonyl]-2-naphthalenyl] oxy]ethyl]-2-naphthalenecarboxamine hydrochloride, was dissolved in DMSO, and cells were treated at a dose of 1µM.

### Neuronal cell cultures

Primary rat striatal cell cultures were generated from E18 striatal tissue as described previously [9,29,30]. Cell culture plates (Denville Scientific, Inc., Plainfield, NJ) were coated overnight with poly-L-lysine (50 µg/mL; Sigma-Aldrich), supplemented with 7.5 µg/mL laminin (Sigma-Aldrich), and rinsed with diH_2_O. Multielectrode arrays (MEAs; Axion Biosystems, Atlanta, GA) were coated with polyethyleneimine (Sigma-Aldrich). Dissected striatal tissue was incubated with papain (LK003178, Worthington Biochemical Corporation, Lakewood, NJ) for 25 min at 37°C. After rinsing in complete Neurobasal media (supplemented with B27 and L-glutamine, Invitrogen), a single-cell suspension was prepared by sequential trituration through large to small fire-polished Pasteur pipettes and filtered through a 100µm cell strainer (Fisher Scientific, Waltham, MA). Cells were pelleted, re-suspended in fresh media, counted, and seeded to a density of 125,000 cells per well on 24-well culture plates (65,000 cells/cm^2^) or 30,000 cells per well on 48-well MEA plates. Cells were grown in complete Neurobasal media for 12 days in vitro (DIV12) in a humidified CO_2_ (5%) incubator at 37°C with half-media changes at DIV1, 4-5, and 8-9. MEAs received a one-half media change to BrainPhys (Stemcell Technologies Inc., Vancouver, Canada) with SM1, L-glutamine supplements, and penicillin-streptomycin starting on DIV4-5 and continued every 3-4 days.

### RNA extraction and RT-qPCR

Total RNA was extracted (RNAeasy kit, Qiagen, Hilden, Germany) and reverse-transcribed (iScript cDNA Synthesis Kit, Bio-Rad, Hercules, CA). cDNA was subject to RT-qPCR for genes of interest, as described previously [9,30]. A list of PCR primer sequences is provided in **Supplemental Table S1**.

### Tissue collection from adult NAc

For RNA sequencing and RT-qPCR experiments, animals were rapidly decapitated either 1hr or 24hr after i.p. injection of either 10 mg/kg cocaine or saline on the final day of testing. Brain tissue was extracted bilaterally and immediately frozen on dry ice. Tissue was stored at −80°C until the day of sequencing or RNA isolation and subsequent RT-qPCR. For CPP experiments, rats were deeply anesthetized with 4-5% isoflurane, then transcardially perfused with formalin (1:10 dilution in PBS, Fisher). Brains were removed and post-fixed for 24hr in formalin at 4°C, protected from light. Fixed tissue was then rinsed and stored in 1xPBS at 4°C until sliced at 50µm using a vibratome. Slices were mounted on glass microscope slides with Prolong Gold anti-fade medium (Invitrogen) containing 4,6-diamidino-2-phenylindole (DAPI) stain as a marker for cell nuclei and coverslipped. Bilateral viral expression and NAc placement were verified manually using a Nikon TiS inverted fluorescent microscope.

### CRISPR/Cas9 and RNAi construct design

CRISPR and shRNA constructs for editing or knockdown of *Gadd45b* were delivered using second-generation lentiviral expression vectors. Cas9 and CRISPR sgRNAs were expressed using a modified version of the lentivirus compatible expression vector lentiCRISPR v2 [31], which was a gift from Feng Zhang (Addgene plasmid #52961; http://n2t.net/addgene:52961; RRID: Addgene_52961). The puromycin resistance cassette in the lentiCRISPR v2 backbone was replaced with a mammalian codon-optimized EGFP tag using MluI and BamHI restriction digest. *Gadd45b*-specific sgRNA targets were designed using online tools provided by the German Cancer Research Center (http://www.e-crisp.org/E-CRISP/). To ensure specificity, CRISPR RNA (crRNA) sequences were analyzed with National Center for Biotechnology Information’s (NCBI) Basic Local Alignment Search Tool (BLAST) to ensure no off-target matches.

Short hairpin RNAs were designed using the Broad Institute Genetic Perturbation Platform web portal. A pLKO.1-TRC vector (a gift from David Root; Addgene plasmid #10879; http://n2t.net/addgene:10879; RRID:Addgene_10879) [32] was cloned into an expression vector with an mCherry reporter (Addgene plasmid #114199) [29] to generate distinct U6-shRNA and EF1α-mCherry expression cassettes. Oligonucleotides containing the shRNA sequence were cloned into this backbone using AgeI and EcoRI restriction digest. A list of the sgRNA and shRNA target sequences is provided in **Supplemental Table S1**.

### Lentivirus production

Viruses were made as described previously [9,29]. All viruses were produced under sterile BSL-2 conditions by transfecting HEK-293T cells (ATCC CRL-3216) with the specified CRISPR or RNAi plasmid, the psPAX2 packaging plasmid, and the pCMV-VSV-G envelope plasmid (Addgene 12260 and 8454) with Fugene HD (Promega, Madison, WI) for 40-48hr in supplemented Ultraculture media (L-glutamine, sodium pyruvate, and sodium bicarbonate) in a T225 culture flask. Supernatant was passed through a 0.45µm filter and centrifuged at 106,883 rcf for 1hr 45 min at 4°C. The viral pellet was resuspended in 1/100^th^ (*in vitro*) or 1/1000^th^ (*in vivo*) supernatant volume of sterile PBS and stored at −80°C. Physical viral titer was determined using either Lenti-X qRT-PCR Titration Kit (Takara, Mountain View, CA) or qPCR Lentivirus Titration Kit (Applied Biological Materials Inc., Richmond, BC, Canada). Viral titers were 1.425 x 10^9^ GC/mL for *lacZ* sgRNA control, 2.663 x 10^9^ GC/mL for *Gadd45b* sgRNA, 2.58 x 10^12^ GC/mL for the *Gadd45b* shRNA, and 7.54 x 10^11^ GC/mL for the scrambled shRNA control. Viruses were stored in sterile PBS at −80°C in single-use aliquots.

### Multi-electrode array recordings

Single-unit electrophysiological activity was recorded using an Axion Maestro Pro recording system (Axion Biosystems). E18 rat primary striatal neurons were seeded in 48-well MEAs at 30,000 cells/well, as described above. Each MEA well within the 48-well plate contains 16 extracellular recording electrodes and a ground electrode. Neurons were transduced with RNAi constructs on DIV5 and MEA recordings were performed on DIV12 while connected to a temperature- and CO_2_-controlled system (maintained at 37°C and 5% CO_2_). Cell health and viral mCherry expression were optically verified using a Nikon TiS inverted fluorescent microscope. Electrical activity was measured by an interface board at 12.5 kHz, digitized, and transmitted to an external computer for data acquisition and analysis in Axion Navigator software (v.1.5.1, Axion Biosystems). All data were filtered using dual 0.01 Hz (high pass) and 5,000 Hz (low pass) Butterworth filters. Action potential thresholds were set automatically using an adaptive threshold for each electrode (> 6 standard deviations from the electrode’s mean signal). Neuronal waveforms collected in Axion Navigator software were exported to Offline Sorter (v.4.0, Plexon, Dallas, TX) for sorting of distinct waveforms corresponding to units on individual electrode channels. Waveforms were sorted by the authors and waveform isolation was confirmed using principal component analysis, inter-spike intervals, and auto- or cross-correlograms. Further analysis of burst activity and firing rate was performed in NeuroExplorer software (v.5.0). Burst activity was analyzed using a poisson burst surprise = 5.

### RNA-sequencing

Bulk RNA-sequencing (RNA-seq) was carried out at the UAB Genomics Core Laboratories at the University of Alabama at Birmingham. RNA was extracted, purified (RNeasy, Qiagen), and DNase-treated for three biological replicates per experimental condition. RNA quality was determined on the BioAnalyzer 2100 (Agilent Technologies, Wilmington, DE). RNA sequencing libraries were created using the NEBNext Ultra II Directional RNA-Seq library kit (NEB, Ipswich, MA) according to manufacturer’s recommendations. The resulting libraries underwent sequencing (75 bp paired-end directional reads; 28.9 - 48.7 million reads*/*sample) on an Illumina NextSeq 500 sequencing platform using standard techniques.

### RNA-seq data analysis

Paired-end FASTQ files were uploaded to the High Performance Computing cluster at the University of Alabama at Birmingham for custom bioinformatics analysis using a pipeline built with snakemake [33] (v5.1.4). Read quality, length, and composition were assessed using FastQC prior to trimming low quality bases (Phred < 20) and Illumina adapters (Trim_Galore! v04.5). Splice-aware alignment to the Rn6 Ensembl genome assembly (v90) was performed with STAR [34] (v2.6.0c). An average of 89.01% of reads were uniquely mapped. Binary alignment map (BAM) files were merged and indexed with Samtools (v1.6). Gene-level counts were generated using the featureCounts [35] function in the Rsubread package (v1.26.1) in R (v3.4.1), with custom options (isGTFAnnotationFile = TRUE, useMetaFeatures = TRUE, isPairedEnd = TRUE, requireBothEndsMapped = TRUE, strandSpecific = 2, and autosort = TRUE). DESeq2 v 1.16.1 (*72*) in R was used to perform count normalization and differential gene expression analysis with the application of Benjamini-Hochberg false discovery rate (FDR) for adjusted p-values. Differentially expressed genes (DEGs) were designated at adjusted *p* < 0.05 and basemean > 50.

Gene enrichment analysis was performed using the Kyoto Encyclopedia of Genes and Genomes (KEGG) network database (version KEGG_09.04.2019) in the ClueGO v2.3.4 application in Cytoscape, using the protein-coding rat genome as a reference set. Enrichment analysis applied Benjamini-Hochberg correction for multiple comparisons and required a minimum of 3 genes per enriched KEGG term category. Significantly enriched categories were designated using a right-sided hypergeometric test, with adjusted *p* < 0.05.

### Reduced representation bisulfite sequencing (RRBS)

RRBS was carried out at the Heflin Center for Genomic Science Genomics Core Laboratories at the University of Alabama at Birmingham. Primary striatal cultures were transduced with scrambled or *Gadd45b* shRNA-expressing lentiviruses at DIV4, and stimulated with 1µM SKF-38393 (or vehicle) for 2hr at DIV11. Genomic DNA from ∼250,000 neurons per sample (*n* = 3 samples per group) was extracted and purified (DNeasy Blood and Tissue DNA extraction kit, Qiagen) prior to RRBS (Ovation RRBS Methyl-Seq System, NuGen, #0353), used according to manufacturer’s instructions. Bisulfite-converted DNA libraries underwent sequencing (75 bp single-end reads; 28.7-35.4 million reads/sample) on an Illumina sequencing platform (NextSeq 500).

### RRBS data analysis

Single-end FASTQ files were uploaded to the High Performance Computing cluster at the University of Alabama at Birmingham for custom bioinformatics analysis using a pipeline built with snakemake [33] (v5.2.2). Read quality, length, and composition were assessed using FastQC prior to trimming low quality bases and kit-specific diversity trimming (Trim_Galore! v04.5). Alignment to the UCSC Rn6 genome assembly was performed with Bismark v0.19.0 (using Bowtie2 v2.3.4.1 and Samtools v1.6). An average of 70.98% of reads were uniquely mapped, and Bismark coverage files generated for CpG methylation were further analyzed in Seqmonk v1.45.4, with unique samples grouped into replicate sets for analysis. DNA methylation status at individual CpGs was quantitated as forward (methylated) reads/total reads at 1,855,525 million CpG sites having > 120 reads for the entire dataset. Differentially methylated CpGs (dmCpGs) across replicate sets were detected using EdgeR [36] (v3.1), setting statistical significance cutoffs at *p* < 0.01 and methylation change > ± 20%.

### Stereotaxic surgery

Naïve adult male Sprague-Dawley rats (Charles River) were anesthetized with 4-5% isoflurane and placed in a stereotaxic apparatus (Kopf instruments, Tujunga, CA). Animals were maintained at a surgical plane of anesthesia with 2-2.5% isoflurane; animals’ respiratory rates were monitored throughout surgery and kept between 35-55 rpm. Surgical coordinates were determined using Paxinos and Watson [37] as a guide to target the NAc core. Under aseptic conditions, guide holes were drilled at AP +1.6 mm, ML ± 1.4 mm, and the infusion needle was lowered to DV −7.0 (all coordinates with respect to bregma). All infusions were made using a gastight 30-gauge stainless steel injection needle and 10µL syringe (Hamilton Company, Reno, NV). Lentivirus constructs were microinfused at a rate of 0.25 µL/min using a syringe pump (Harvard Apparatus, Holliston, MA), totaling 2 µL per hemisphere. Following each infusion, needles remained in place for 10 min to allow for diffusion of virus. After retracting the infusion needles, guide holes were filled with sterile bone wax and the surgical incision was closed with nylon sutures. Animals received buprenorphine (0.03 mg/kg) and carprofen (5 mg/kg) for pain management and inflammation, as well as topical bacitracin (500 units) to prevent infection at the incision site.

### Acute cocaine locomotor testing and cocaine sensitization

For open-field locomotor testing, naïve male animals (*n* = 12 per group) were given i.p. injections of saline on days 1 and 2 immediately before 30 min of locomotor testing. On days 3 and 10, half of the animals were given an i.p. injection of 10 mg/kg cocaine, and the other half received i.p. injections of saline. Locomotor activity was monitored in a 43cm x 43cm plexiglass locomotor activity chamber (Med Associates, Inc., St. Albans, VT) with opaque white wall covering and an open top. Each chamber included a 48-channel X-Y infrared array (Med Associates, Inc.) that was used to measure distance traveled in conjunction with Activity Monitor software (Med Associates, Inc.). Test chambers were cleaned with 0.0156% chlorhexidine and 70% ethanol at the beginning and end of each testing day, and cleaned with 70% ethanol in between trials. Animals were transported to the behavioral testing core 30 min prior to testing on a covered cart. Behavioral test sessions were conducted during the light cycle, and the overhead lights and white noise generator remained on in the testing room. The same male experimenter conducted all locomotor testing and was present in the room during testing.

### Conditioned place preference testing

Conditioned place preference (CPP) testing was completed in a three-chamber apparatus with guillotine doors (Med Associates, Inc.). The two chambers used for conditioning measured approximately 27cm x 21cm x 22cm, one with opaque black walls and stainless steel bar flooring, and the other with opaque white walls and metal wire grid flooring. The conditioning chambers were separated from one another by guillotine-style doors and a central grey compartment with solid flooring, measuring 12cm x 21cm x 22cm. All three chambers had a clear perforated acrylic lids with house lights. Each apparatus included a 16-channel infrared controller (Med Associates, Inc.) to track rodent position, in conjunction with Med PC software (v4.1.49; Med Associates, Inc.).

For CPP testing in transgenic *Gadd45b* knockout mice, naïve male animals (*n* = 7-9 per group) were placed in the central compartment on the first day of testing (i.e. pre-test) and were permitted to explore all 3 chambers of the CPP apparatus during a 20 min session. On days 2 and 4, mice were given an i.p. injection of saline immediately prior to being placed in the initially preferred chamber for the 20 min conditioning session. On days 3 and 5, mice were given an i.p. injection of 10 mg/kg cocaine before being placed in the initially non-preferred chamber for the 20 min conditioning session. On day 6 (i.e. post-test), mice were again placed in the central compartment and allowed to freely move between all three chambers. Time spent in each chamber during the post-test was calculated and compared to the pre-test times. Overhead lights were off, and CPP house lights were set to an intensity of 10. Cedar and pine beddings were used in the white and black chambers, respectively, to help mice further differentiate between the environments.

CPP testing in adult male rats (*n* = 7-8 per group) began 2 weeks following viral infusion surgeries. Days 1-6 of testing were identical to CPP testing in transgenic mice. However, this schedule was repeated in rats on days 7-10, increasing the cocaine dose to 20 mg/kg for cocaine conditioning on days 8 and 10. On day 11, rats underwent a second post-test to measure cocaine place preference. Overhead lights were left on, and CPP house lights were set to an intensity of 7-8. Testing and conditioning sessions lasted 30 min.

### Statistical analysis

Sample sizes were calculated using a freely available calculator [38]. Transcriptional differences from RT-qPCR experiments were compared with an unpaired t-test with Welch’s correction, one-way ANOVA with Tukey’s *post hoc* tests, or two-way ANOVA with Sidak’s *post hoc* tests, where appropriate. MEA data was compared with Mann-Whitney U-tests. Locomotor sensitization data was compared using a two-way ANOVA with Tukey’s *post hoc* tests where appropriate. CPP data was compared with a two-way ANOVA with Bonferroni’s multiple comparisons test. Statistical significance was designated at α = 0.05 for all analyses. Statistical and graphical analyses were performed with Prism software (GraphPad, La Jolla, CA). Statistical assumptions (e.g. normality and homogeneity for parametric tests) were formally tested and examined via boxplots.

### Data availability

Sequencing data that support the findings of this study will be deposited at Gene Expression Omnibus. All relevant data that support the findings of this study are available by request from the corresponding author (J.J.D.).

## Results

To examine the effects of cocaine on *Gadd45b* expression *in vivo*, we collected NAc tissue of adult naïve male rats at 1hr and 24hr following treatment with either cocaine (10 mg/kg, i.p.) or saline. As expected, mRNA for several classic IEGs (*Arc, Egr1, Fos, Fosb*, and Δ*Fosb*) was transiently increased 1hr after cocaine injection, with a return to baseline at 24hr (**Figure 1a**). Similarly, RT-qPCR for *Gadd45b* revealed significant upregulation at 1hr following cocaine treatment, but not at 24hr (**Figure 1b**). Given that GADD45 proteins have been implicated in active demethylation [23,24], we also measured expression of other genes involved in DNA methylation, such as *Dnmt3a, Dnmt3a1, Dnmt3a2, Dnmt3b, Tet1*, and *Tet3* (**Figure 1b**). However, expression of these genes did not differ between cocaine- and saline-treated animals at either timepoint.

**Figure 1.**
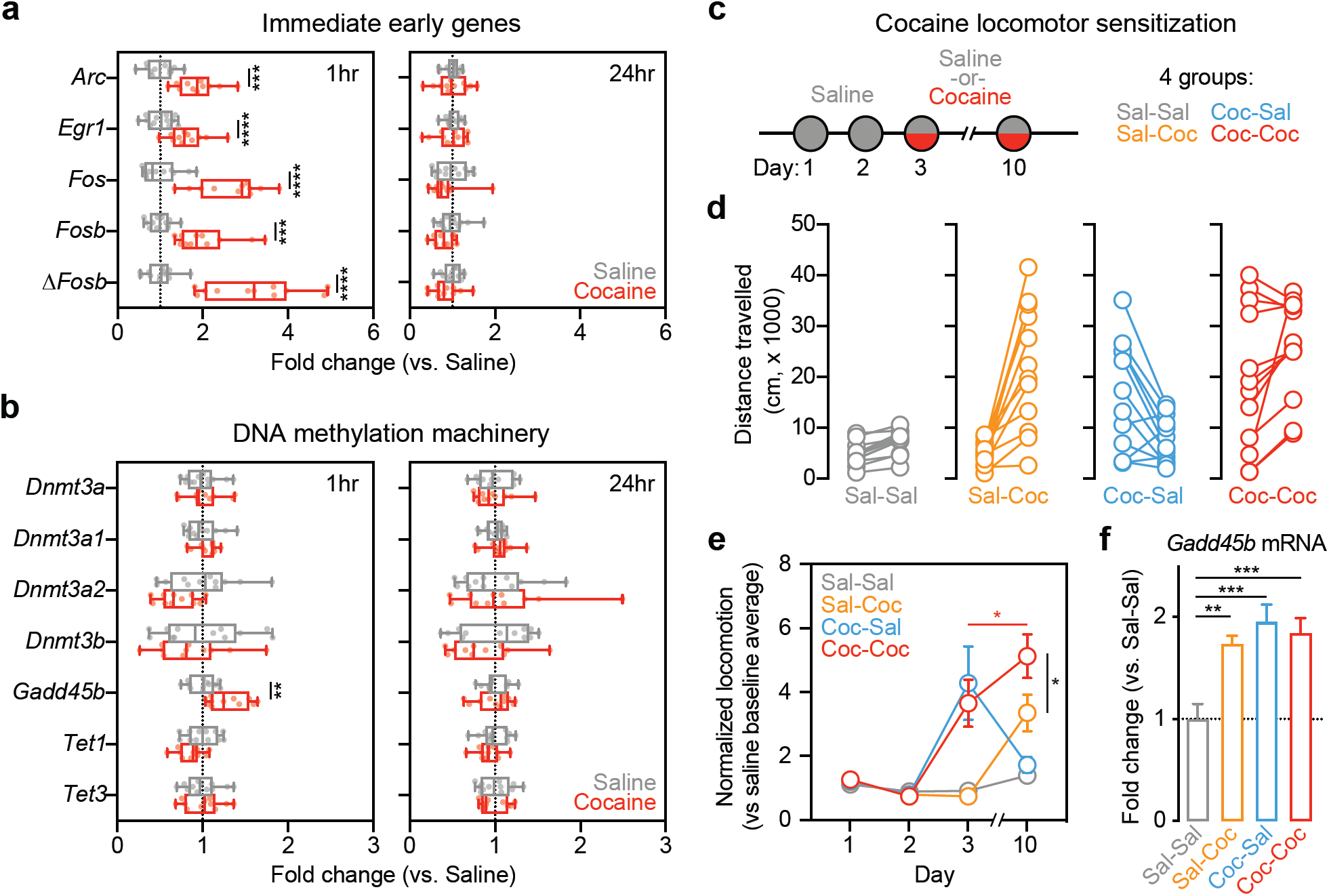
Exposure to acute cocaine leads to increased expression of immediate early genes and *Gadd45b in vivo*. **a**, Adult male rats were administered a single i.p. injection of saline (*n* = 12) or 10 mg/kg cocaine (*n* = 11) and returned to their home cage. NAc tissue was collected bilaterally at either 1hr or 24hr post-injection. RT-qPCR reveals significant increases in immediate early gene (IEG) expression at 1hr in cocaine-treated rats compared to saline controls (multiple *t*-tests, *Arc t*_(21)_ = 5.188, adjusted *p* = 0.000116, *Egr1 t*_(21)_ = 4.078, adjusted *p* = 0.000539, *Fos t*_(21)_ = 6.552, adjusted *p* = 0.000008, *Fosb t*_(21)_ = 4.919, adjusted *p* = 0.000145, Δ*Fosb t*_(21)_ = 6.581, adjusted *p* = 0.000008). No changes were observed at 24hr. **b**, When examining several genes involved in activity-dependent DNA methylation and demethylation processes, only *Gadd45b* mRNA was significantly increased 1hr following acute exposure to cocaine (multiple *t*-tests, *Gadd45b t*_(21)_ = 3.651, adjusted *p* = 0.010389). There were no significant changes observed at 24hr. **c**, To examine cocaine locomotor sensitization, all rats were administered saline on days 1 and 2. On days 3 and 10, half received a 10 mg/kg cocaine injection. This yeilded four groups (*n* = 12/group). **d**, Distance travelled was significantly increased in animals that received cocaine either on day 3 or on day 10. Animals which received two consecutive cocaine injections on day 3 and day 10 also exhibited increases in locomotor activity. **e**, Locomotor sensitization was only observed in animals that received cocaine on both day 3 and day 10 (two-way ANOVA, *F*_(9,132)_ = 9.714, *p* < 0.0001 for day x group interaction; Tukey’s *post hoc* tests, *p* < 0.05 for indicated comparisons). **f**, NAc tissue was collected bilaterally 1hr post-injection on day 10 for RT-qPCR analysis. *Gadd45b* mRNA was significantly increased in all groups with cocaine experience (one-way ANOVA, *F*_(3,44)_ = 8.9925, *R*^2^ = 0.38, *p* < 0.0001; Tukey’s *post hoc* tests, \*\**p* < 0.01 and \*\*\**p* < 0.001 for indicated comparisons). All data are expressed as mean ± s.e.m.

In order to better characterize how cocaine experience affects *Gadd45b* expression, we used a locomotor sensitization paradigm [39] in which animals received an injection of saline on days 1-2 and an injection of either saline or cocaine (10 mg/kg) on days 3 and/or 10 (**Figure 1c**). Acutely, animals that received cocaine displayed heightened locomotor activity compared to saline controls, and locomotor sensitization was only observed in animals that received cocaine on both day 3 and day 10 (**Figure 1d-e**). NAc tissue was collected 1hr following treatment on day 10, and RT-qPCR revealed that *Gadd45b* mRNA was upregulated in all groups with cocaine experience (**Figure 1f**), suggesting that *Gadd45b* is induced by cocaine and cocaine-paired environments, independent of the acute psychostimulant action of the drug.

Given that *Gadd45b* is important for learning and memory [27,40] and is upregulated following acute cocaine experience, we next aimed to determine if *Gadd45b* expression is necessary for cocaine reward memory. To do this, we employed a CPP paradigm, which is commonly used to examine drug-associated memories and the rewarding properties of drugs of abuse. In this assay, drug-naïve animals freely explore a three-chambered apparatus on day 1 (**Figure 2b**) to assess initial preference for a particular context. Following four days of conditioning, animals are again allowed to freely explore the apparatus on day 7 in the absence of the drug to determine if they exhibit a preference for the drug-paired context. Following conditioning, wild-type mice formed a normal CPP response for the cocaine-paired chamber. However, this cocaine-paired place preference was attenuated in Gadd45b knockout mice (**Figure 2a-c**) despite similar time spent on the cocaine-paired side during the pre-test, suggesting a deficit in cocaine reward memory.

**Figure 2.**
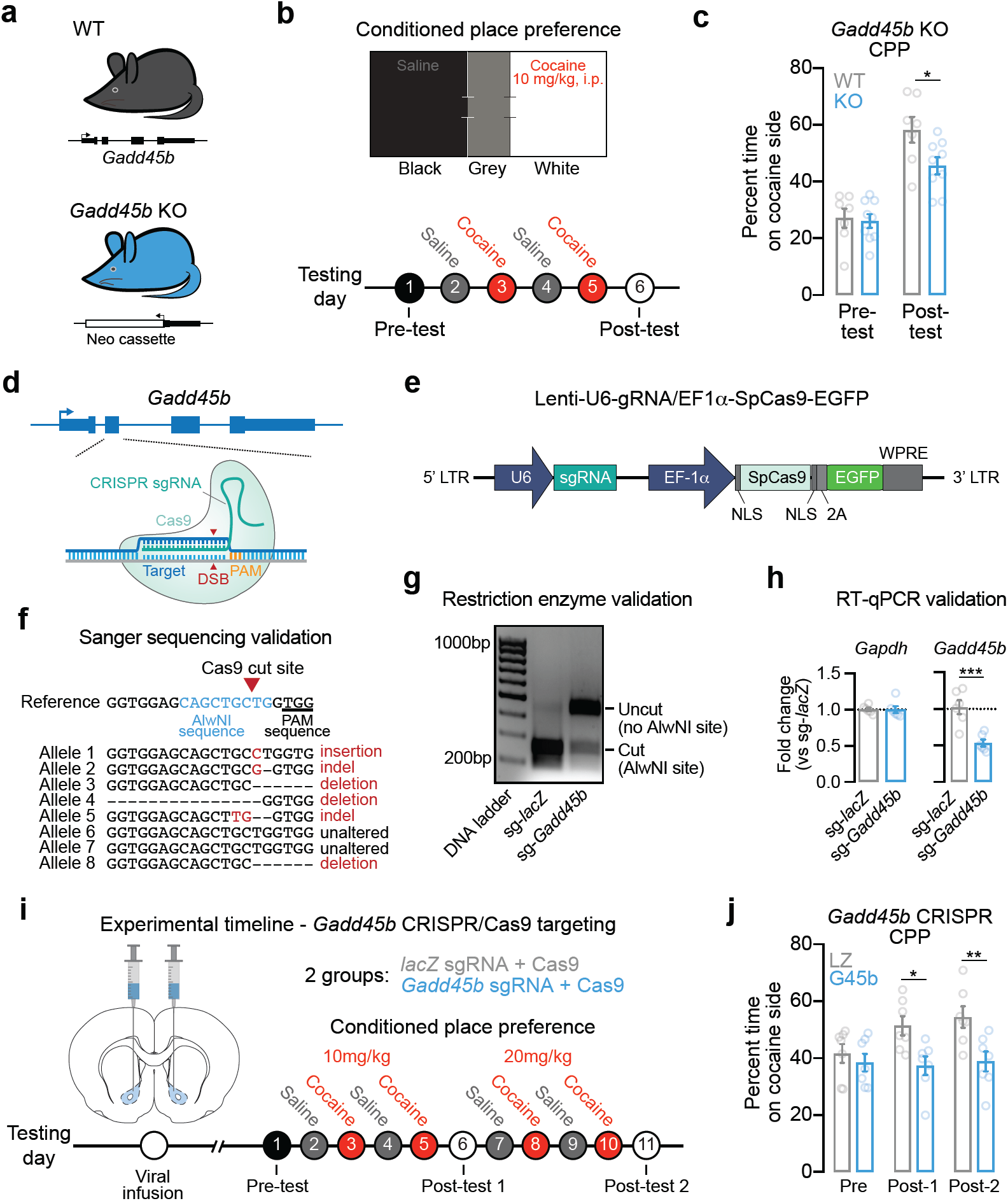
Efficient gene knockdown of *Gadd45b* attenuates cocaine reward memory. **a**, Male wild-type (*n* = 7) and mutant *Gadd45b* knockout (*n* = 9) mice between 2-6 months of age were used for behavioral testing. **b**, Conditioned place preference (CPP) apparatus and experimental timeline for CPP testing. **c**, Cocaine-paired place preference was significantly attenuated in *Gadd45b* mutant mice (two-way ANOVA, *F*(1,14) = 6.232, *p =* 0.0256 for group comparison; Bonferroni’s *post hoc* multiple comparisons, *p* < 0.05). **d**, Targeting strategy to induce insertion/deletion mutations in *Gadd45b*. CRISPR guide RNAs (gRNAs) were engineered to target exon 2 of *Gadd45b*. **e**, Illustration of lentivirus construct designed to express gRNA (driven by the U6 promoter) and the Cas9 nuclease (driven by the Ef1a promoter), with bicistronic expression of EGFP. **f**, Sanger sequencing of individual genomic DNA clones reveals CRISPR-mediated mutation in 6 of 8 alleles (75% efficiency) following lentiviral transduction. Neurons were transduced with lentivirus at DIV7 and genomic DNA was extracted at DIV11. **g**, Genomic DNA restriction digest at AlwNI site (which is mutated by Cas9 editing) reveals almost complete digestion in *lacZ* gRNA control, but loss of digestion when AlwNI site has been mutated in *Gadd45b* gRNA group. **h**, RT-qPCR validation demonstrates robust knockdown of *Gadd45b* mRNA (*n* = 6 per group; unpaired *t*-test, *t*(10) = 4.615, *R*^2^ = 0.68, *p* = 0.001). Reminaing mRNA contains insertions/deletions resulting in premature translation termination or dysfunctional protein. **i**, Experimental timeline for CRISPR/Cas9 viral transduction and cocaine CPP testing. Lentiviral constructs targeting *lacZ* (*n* = 8) or *Gadd45b* (*n* = 7) were bilaterally injected into the NAc core 2 wks prior to behavioral testing. **j**, Rats that received the *Gadd45b*-targeted CRISPR/Cas9 construct failed to form a place preference to cocaine at both 10 mg/kg and 20 mg/kg (two-way ANOVA, *F*(1,12) = 8.053, *p* = 0.0150 for group comparison; Bonferroni’s *post hoc* multiple comparisons, \**p* < 0.05 and \*\**p* < 0.01 for indicated comparisons). All data are expressed as mean ± s.e.m.

We next sought to confirm this effect in the absence of possible developmental compensatory mechanisms in a germline mutant model. Additionally, we sought to better characterize the specific role of *Gadd45b* in the ventral striatum. We repeated this behavioral assay using *Gadd45b*-targeted CRISPR/Cas9 gene editing in the ventral striatum in rats in order to produce site-specific insertion/deletion events that render dysfunctional GADD45B protein. We designed lentiviral constructs expressing the Cas9 nuclease and CRISPR single guide RNA (sgRNA) targeted to exon 2 of *Gadd45b* (**Figure 2d-e**) and validated successful gene editing *in vitro* using three complementary approaches. Sanger sequencing revealed *Gadd45b* locus mutations in 6 out of 8 alleles following transduction with CRISPR constructs, indicating 75% editing efficiency (**Figure 2f**). Furthermore, restriction enzyme digest of genomic DNA at an AlwNI site located within the sgRNA sequence (overlapping the Cas9 cut site) revealed near-complete digestion in the *lacZ* non-targeting sgRNA control, indicative of an intact AlwNI site. In contrast, the *Gadd45b* sgRNA group lacked AlwNI-mediated digestion, confirming Cas9-driven mutation at the AlwNI site (**Figure 2g**). As a final validation measure, RT-qPCR for *Gadd45b* mRNA indicated significant reduction in *Gadd45b* sgRNA-targeted cells (as compared to the *lacZ* sgRNA control), demonstrating that gene editing was sufficient to reduce total levels of *Gadd45b* expression (**Figure 2h**). Adult male rats (*n* = 7-8 per group) underwent stereotaxic surgery to infuse the CRISPR/Cas9 + sgRNA lentivirus bilaterally targeting the NAc core. After allowing two weeks for viral expression, CPP testing began (**Figure 2i**). Whereas both groups exhibited a similar preference prior to cocaine pairing, only the *lacZ*-targeted animals developed a place preference for cocaine at the two doses tested (**Figure 2j**). Rats that received the *Gadd45b*-targeted CRISPR/Cas9 gene editing constructs failed to develop a cocaine-paired place preference at both 10mg/kg and 20mg/kg, suggesting that *Gadd45b* in the NAc is necessary for cocaine reward memory.

In order to further examine how DA regulates *Gadd45b* expression, we first used a well-established and highly controllable rat primary striatal neuron culture system [9,29,30] to examine DA-dependent *Gadd45b* induction (**Figure 3a**). DIV11 striatal cultures were treated with 1µM DA for 1hr, a treatment which closely models both the increase in DA concentration and the temporal dynamics of DA changes in the striatum *in vivo* following acute cocaine exposure [6,7,41]. Using a recently published dataset in which RNA-seq was performed on striatal neurons 1hr after DA treatment [9], we found that *Gadd45b* was identified as one of only 100 mRNAs increased by DA (**Figure 3b**). To validate this finding, as well as dissect the relevant DA receptor systems integral to *Gadd45b* action, we performed RT-qPCR on DA-treated striatal cultures (**Figure 3a**). In agreement with RNA-seq results, we found that acute DA treatment significantly increased *Gadd45b* mRNA (**Figure 3c**). DA-induced increases in *Gadd45b* mRNA were blocked in the presence of the DRD1 antagonist SCH-23390 (1µM; **Figure 3d**). Similarly, treatment with DRD1 agonist SKF-38393 (1µM), but not DRD2/DRD3 agonist Quinpirole (1µM), significantly increased *Gadd45b* expression compared to vehicle-treated controls (**Figure 3e**). *Gadd45b* mRNA was also elevated following stimulation with Forskolin (20µM), an activator of adenylyl cyclase that mimics DRD1-mediated G-protein coupled receptor signaling pathways (**Figure 3f**). In order to better characterize the signaling cascades necessary for DA-induced increases in *Gadd45b*, we next co-treated striatal cultures with DA and either a CREB inhibitor (666-15, 1µM) or a MEK inhibitor (U0126, 1µM). CREB inhibition decreased *Gadd45b* mRNA both at baseline and following DA treatment (**Figure 3g**). Similarly, MEK inhibition blocked baseline and DA-induced *Gadd45b* expression, compared to vehicle- or U0124-treated controls (**Figure 3h**). Together, these data demonstrate that DA-mediated induction of *Gadd45b* requires DRD1 signaling through MAPK and CREB signaling pathways.

**Figure 3.**
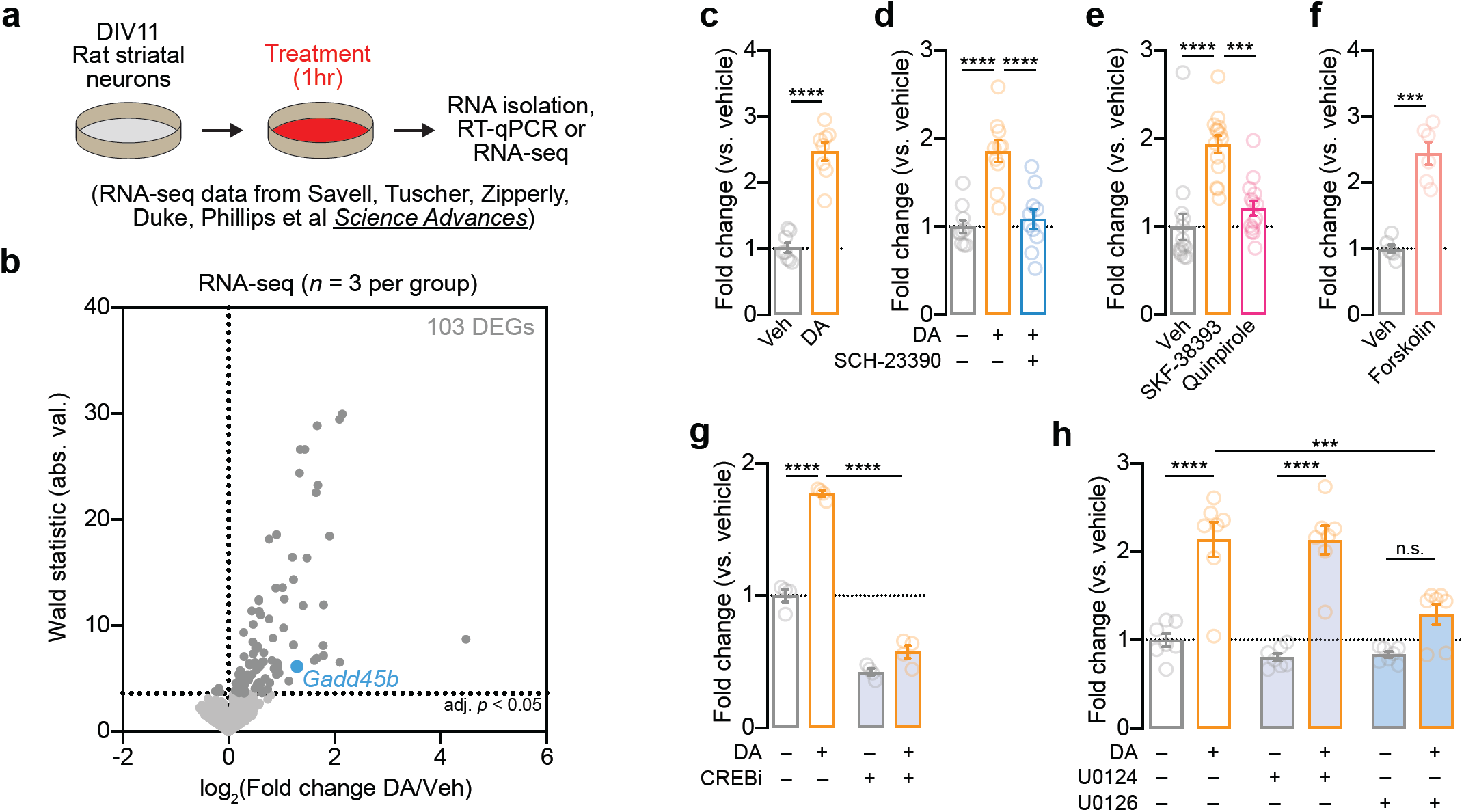
Induction of *Gadd45b* mRNA by dopamine requires DRD1 dopamine receptors and CREB activation *in vitro*. **a**, Illustration of experimental design using primary rat striatal culture system. **b**, Unbiased identification of DA-induced DEGs with RNA-seq reveals *Gadd45b* as one of 103 DA-induced IEGs. Cells were treated with DA (1μM) for 1hr prior to RNA isolation and RNA-seq. **c**, RT-qPCR validation of DA-induced increase in *Gadd45b* mRNA (*n* = 8 per group; unpaired *t*-test, *t*_(14)_ = 9.377, *R*^2^ = 0.86, *p* < 0.0001). **d**, DA-induced increases in *Gadd45b* mRNA are blocked by co-treatment with the DRD1 receptor antagonist SCH-23390 (1μM; *n* = 10 per group; one-way ANOVA, *F*_(2,27)_ = 20.78, *R*^2^ = 0.61, *p* < 0.0001; Tukey’s *post hoc* test, *p* < 0.0001 for indicated comparisons). **e**, *Gadd45b* mRNA induction is mimicked by DRD1 receptor agonist SKF-38393 (1μM), but not the D2/D3 receptor agonist Quinpirole (1μM; *n* = 14 per group; one-way ANOVA, *F*_(2,39)_ = 18.73, *R*^2^ = 0.49, *p* < 0.0001; Tukey’s post-hoc test, *p* < 0.001 for indicated comparisons). **f**, *Gadd45b* mRNA is induced by the adenylyl cyclase activator Forskolin (20μM; *n* = 6 per group) unpaired *t*-test with Welch’s correction, *t*_(6.198)_ = 7.781, *R*^2^ = 0.91, *p* = 0.0002). **g**, CREB inhibition with 666-15 (1μM) blocks baseline and DA-induced increases in *Gadd45b* mRNA (*n* = 4 per group; two-way ANOVA, *F*_(1,12)_ = 560.3, *p* < 0.0001 for main effect of CREB inhibition; Sidak’s *post hoc* tests, *p* < 0.0001 for indicated comparisons). **h**, MEK inhibition with U0126 (1μM) prevents DA-induced increases in *Gadd45b* mRNA (*n* = 7 per group; two-way ANOVA, *F*_*(2, 36)*_ = 26.51, *p* = 0.0004 for main effect of MEK inhibition; Sidak’s *post hoc* tests, \*\*\*\**p* < 0.0001 and \*\*\**p* = 0.0003 for indicated comparisons). U0124 (1μM) is an inactive analog of U0126 and was included as a negative control.

*Gadd45b* is involved in many cellular processes, including those related to learning and memory, neurodevelopment, and cellular stress [27,40,42–45]. To further explore the molecular roles of *Gadd45b in vitro*, we designed a custom short-hairpin RNA (shRNA) interference approach to reduce *Gadd45b* mRNA. Lentiviral transduction with *Gadd45b* shRNA produced > 90% knockdown of *Gadd45b* mRNA without alterations in mRNA from the control housekeeping gene *Gapdh* (**Figure S1**). Bulk RNA-seq of striatal cell cultures identified 7,325 DEGs following *Gadd45b* knockdown, consisting of 3,527 downregulated genes and 3,798 upregulated genes (**Figure 4a-c; Table S2**). KEGG network analysis revealed that genes involved in addiction, as well as dopamine and glutamatergic synapses, were significantly downregulated following *Gadd45b* knockdown (**Figure 4d**). In order to characterize which gene programs are active following DRD1 activation, we treated striatal neurons with 1µM SKF-38393. Compared to scrambled controls, neurons transduced with *Gadd45b* shRNA exhibited a dampened transcriptional response (**Figure 4e; Table S3**). Interestingly, many of the genes that are most highly upregulated following DRD1 stimulation in control neurons were blunted in SKF-treated neurons transduced with *Gadd45b* shRNA (**Figure 4e-g**). These findings suggest that *Gadd45b* knockdown results in extensive transcriptional dysregulation and prevents the full activation of DA-responsive gene programs in striatal neurons.

**Figure 4.**
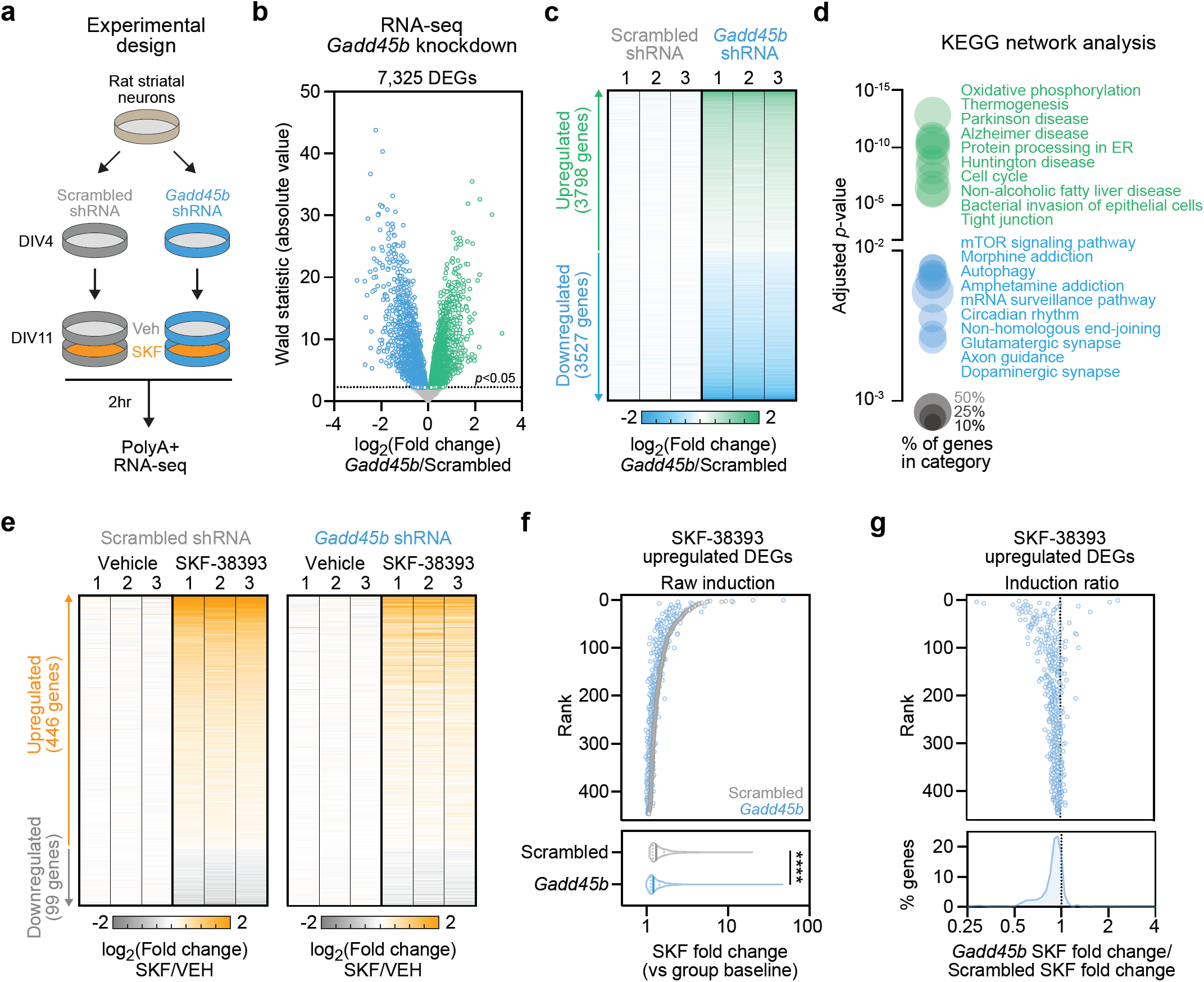
*Gadd45b* knockdown causes widespread transcriptional dysregulation and blocks dopamine-dependent gene programs in striatal neurons. **a**, Illustration of experimental design for shRNA-mediated *Gadd45b* knockdown and transcriptional profiling. **b**, Volcano plot of 7,325 differentially expressed genes (DEGs; adjusted *p* < 0.05) in *Gadd45b* shRNA group as compared to scrambled shRNA control group. **c**, Heatmap showing replicate values of all DEGs (3,798 upregulated, 3,527 downregulated) after *Gadd45b* knockdown. **d**, KEGG network analysis reveals differential upregulation of genes implicated in oxidative phosphorylation and neurodegenerative disease categories, and downregulation of genes implicated in dopaminergic synapse function and addiction categories. Adjusted *p* < 0.05 for all categories shown. **e**, Heatmap of DEGs altered by 2hr stimulation with the DRD1 receptor agonist SKF-38393 (1µM), stratified by shRNA treatment group. **f-g**, Genes upregulated by SKF-38393 stimulation are significantly less induced under conditions of *Gadd45b* knockdown (ratio paired *t*-test for scrambled vs *Gadd45b* shRNA comparison of SKF-38393 fold change, *t*_(445)_ = 13.57, *p* < 0.0001, *R*^*2*^ = 0.29).

Given that *Gadd45b* has previously been implicated in DNA methylation dynamics in the nervous system [19,20,46–48], we next explored whether *Gadd45b* knockdown altered DNA methylation in striatal neuron cultures. Using reduced representation bisulfite sequencing (RRBS), we profiled ∼1.86 million unique CpG sites following *Gadd45b* shRNA (or scrambled shRNA control) in combination with DRD1 receptor stimulation (vehicle or 1µM SKF-38393 treatment; **Figure S2a**). As expected, DNA methylation patterns exhibited typical bimodal distribution of CpG methylation, with depletion at CpG islands and gene promoters and CpG methylation enrichment within gene bodies (**Figure S2b-e**). Surprisingly, *Gadd45b* shRNA did not alter global DNA methylation profiles, as cells with *Gadd45b* knockdown displayed nearly identical DNA methylation levels at regulatory elements, promoters, and gene bodies (**Figure S2c-e**). Further, only 129 CpGs (0.006% of CpGs) were differentially methylated (termed dmCpGs) after *Gadd45b* knockdown (**Table S4**), suggesting largely intact global and site-specific DNA methylation landscapes.

To examine stimulus-induced changes in DNA methylation, we quantified SKF-38393-dependent changes in DNA methylation in the scrambled shRNA control group, and identified 1930 dmCpGs following SKF-38393 treatment as compared to vehicle-treated controls (**Figure S2f; Table S5**). Notably, SKF-38393-mediated changes in CpG methylation were absent following *Gadd45b* knockdown (**Figure S2f-j**), suggesting that *Gadd45b* is necessary for Drd1-dependent changes in DNA methylation. Surprisingly, while *Gadd45b* has been demonstrated to contribute largely to activity-induced DNA hypomethylation, *Gadd45b* shRNA produced robust deficits in both CpG hypermethylation and hypomethylation events following SKF-38393 exposure (**Figure S2h-j**). Taken together with RNA-seq results, these findings demonstrate that *Gadd45b* is required for key transcriptional and epigenetic alterations downstream of DA receptor activation.

Given the large-scale downregulation of genes related to glutamatergic and dopaminergic synaptic function following *Gadd45b* knockdown (**Figure 4d**), we next sought to determine whether *Gadd45b* manipulation alters physiological properties of striatal neurons. We transduced striatal cell cultures with shRNA lentiviral constructs prior to electrophysiological characterization using a high-throughput multielectrode array system (**Figure 5a-d**). Following transduction on DIV5, we verified expression of shRNA constructs by visualizing mCherry expression (**Figure 5b**). On DIV12, we performed a 20 min baseline extracellular electrophysiological recording, then treated cells with either vehicle or SKF-38393 (1µM) and recorded activity for an additional 1hr. At baseline, neurons transduced with *Gadd45b* shRNA did not differ from scrambled controls in the number of spontaneously active units (*n* = 996-1089 units across 36 wells/group), mean firing rate, mean action potential burst frequency, or percent of spikes occurring in a burst (**Figure 5e-h**). However, we found a significant reduction in burst duration in *Gadd45b* knockdown neurons as compared to scrambled controls (**Figure 5i**). Application of DRD1 agonist SKF-38393 increased action potential firing rate in both scrambled and *Gadd45b* shRNA neurons, compared to vehicle-treated wells (**Figure 5j-l**). When examining SKF-38393 response as a function of baseline action potential firing rate, we found no difference in SKF-38393 response between the *Gadd45b* knockdown group and scrambled controls, and this lack of effect was maintained across all observed baseline frequencies (**Figure 5m**). Thus, *Gadd45b* knockdown significantly reduces burst duration in striatal neurons, without affecting other baseline electrophysiological characteristics or response to DRD1 receptor activation.

**Figure 5.**
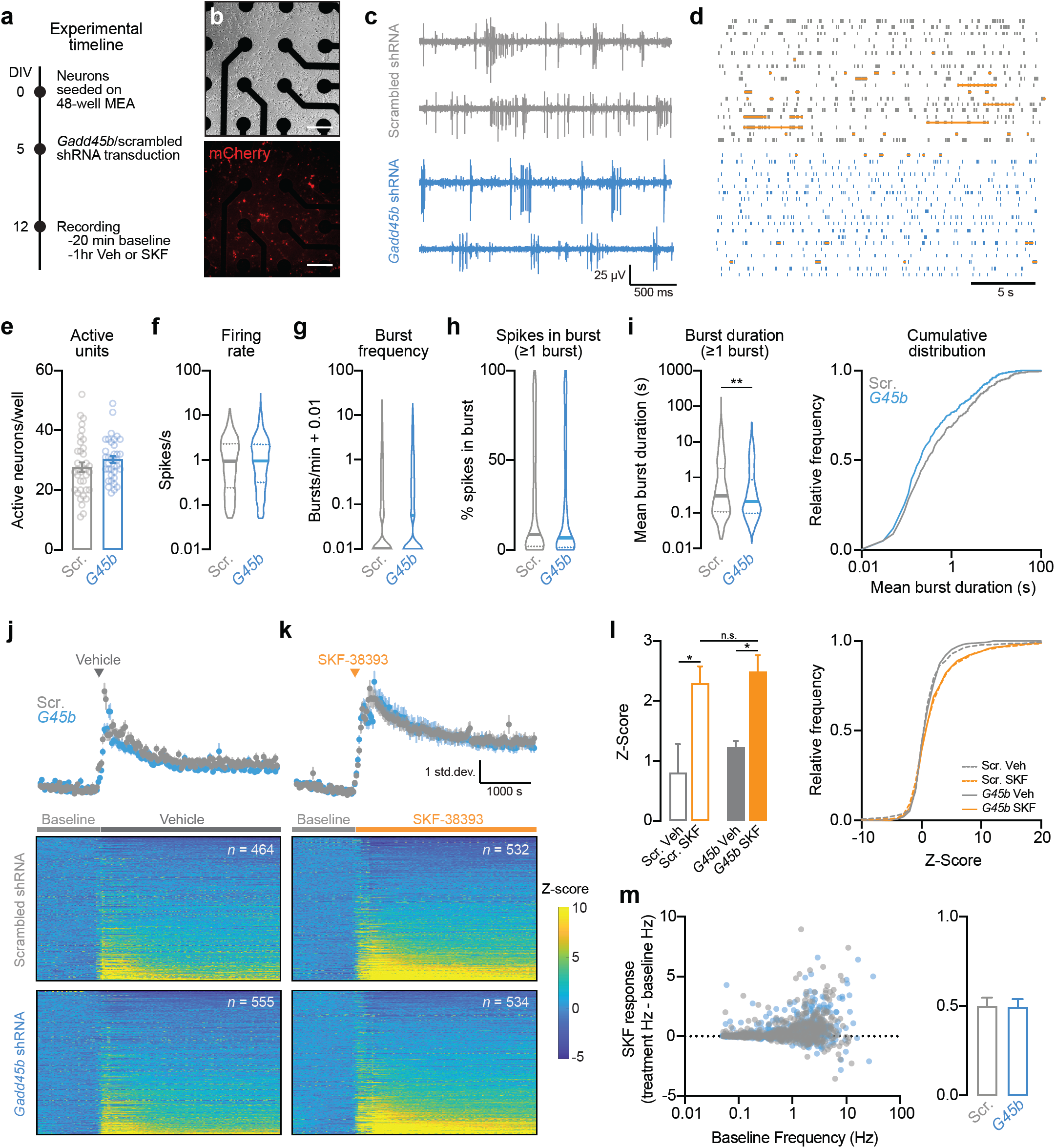
shRNA knockdown of *Gadd45b* in primary striatal neurons alters burst duration, without affecting other electrophysiological characteristics or response to DRD1 stimulation. **a**, Experimental timeline for viral transduction and extracellular single-unit recordings. Primary striatal neurons were grown on multielectrode arrays (MEAs) and transduced with lentiviral shRNA constructs on DIV5. **b**, Live cell imaging of transduced neurons (expressing mCherry) on DIV13 after collecting electrophysiological measures. Scale bar = 100 μm. **c**, Representative traces of four units transduced with either scrambled shRNA or *Gadd45b* shRNA. **d**, Representative raster plots (one neuron per row). Orange horizontal lines denote action potential bursts. **e**, *Gadd45b* knockdown does not affect the number of spontaneously active units per well (*n* = 36 wells/group). **f-g**, Action potential frequency and burst frequency are not different between neurons transduced with *Gadd45b* shRNA (*n* = 1089 units) and scrambled controls (*n* = 996 units). **h**, For all neurons with at least one burst, *Gadd45b* knockdown does not alter % of spikes occuring in a burst. **i**, shRNA knockdown of *Gadd45b* (*n* = 504) significantly decreases burst duration (Mann-Whitney *U* test, *U* = 105010, *p* = 0.0052) compared to scrambled controls (*n* = 465). **j-k**, Mean Z-score (top) and individual neuron heatmap (bottom) following application of vehicle or 1µM SKF-38393. **l**, SKF-38393 increases neuronal firing rate as compared to vehicle control (One-way Kruskal-Wallace ANOVA, *F* = 12.43, *p* = 0.0061; Dunn’s multiple comparisons test, \**p* < 0.05). Effects of SKF-38393 did not differ based on *Gadd45b* knockdown status. **m**, Left, SKF-38393 response as a function of baseline (pre-SKF-38393) action potential frequency. Right, SKF-38393 response did not differ in *Gadd45b* shRNA treatment group. All bar graphs expressed as mean ± s.e.m. All violin plots expressed as median ± quartiles.

## Discussion

Here, we present evidence that *Gadd45b* is an IEG induced in striatal neurons following acute cocaine exposure *in vivo* and DA treatment *in vitro*. Furthermore, *Gadd45b* mRNA is upregulated following exposure to cocaine-paired contexts and is necessary for cocaine CPP. In cultured striatal neurons, we demonstrate that *Gadd45b* induction requires DRD1 activation, as well as MAPK and CREB signaling cascades. Knockdown of *Gadd45b* in striatal neurons blocks the expression of DRD1-responsive genes and decreases the expression of genes implicated in addiction and dopaminergic signaling. Additionally, DRD1-mediated bidirectional DNA methylation changes are also inhibited by *Gadd45b* knockdown, irrespective of initial methylation status. Finally, we demonstrate that *Gadd45b* knockdown significantly decreases the duration of action potential bursts in striatal neurons, without altering other electrophysiological characteristics. Interestingly, while *Gadd45b* knockdown blocks DRD1-mediated changes in gene expression and DNA methylation, medium spiny neurons are able to maintain increased action potential firing following DRD1 activation regardless of *Gadd45b* knockdown status.

Originally termed *MyD118, Gadd45b* was first characterized as a primary response gene important in the myeloid differentiation program [42,43,49]. *Gadd45b* is broadly induced by a variety of genotoxic and environmental stimuli, including DNA alkylating agents [43–45], UV radiation [43–45,50], anisomycin treatment [43], and exposure to a hyperosmotic environment [43]. In the hippocampus, *Gadd45b* mRNA is increased following contextual fear conditioning [40], electroconvulsive seizure [21,51], spatial exploration of a novel environment [21], exercise [21], and kainic acid treatment [52–54]. Furthermore, activity-dependent *Gadd45b* induction in hippocampal neurons is dependent on the *N*-methyl-D-aspartate receptor (NMDAR), calcium, and calcium/ calmodulin-dependent protein kinase signaling, suggesting that *Gadd45b* is induced in the same manner as classic IEGs, such as *Arc* [21]. Additionally, seizure-induced increases in *Gadd45b* are CREB-dependent [52]. *Gadd45b* has also been shown to act as an IEG in other brain regions. For instance, increases in *Gadd45b* mRNA are observed in the NAc after both optical stimulation of VTA DA neuron terminals in the NAc and following direct infusion of brain-derived neurotrophic factor (BDNF) into the NAc [22]. Here, we demonstrate that *Gadd45b* acts as an IEG in the ventral striatum following cocaine administration *in vivo* (**Figure 1b**,**f**) and DRD1 activation *in vitro* (**Figure 3c-e**), and we show for the first time that striatal induction of *Gadd45b* is dependent on both MAPK and CREB signaling (**Figure 3g-h**). Furthermore, we demonstrate that striatal neurons are still capable of physiological responses to DRD1 stimulation following *Gadd45b* knockdown (**Figure 5j-m**). This finding suggests that the primary deficit in neurons lacking *Gadd45b* is the failure to translate neuronal activity into a programmed transcriptional and epigenetic response. In addition, the relative lack of change in many neuronal physiological properties following *Gadd45b* manipulation (**Figure 5e-h**) indicates that *Gadd45b*-mediated transcriptional reorganization was not associated with overt changes in neuronal phenotype, altered neuronal health, or overall cell death.

While *Gadd45b* has been previously investigated in the context of learning and memory, its role in these processes remains unclear. Increased *Gadd45b* expression has been observed following the induction of long-term potentiation (LTP), a cellular correlate of learning and memory, alongside other genes with well-established roles in neural plasticity [53,54]. However, following a near-threshold stimulus, hippocampus slices from *Gadd45b*^*-/-*^ mutant mice exhibit enhanced late-phase LTP, suggesting a plasticity-repressive role of *Gadd45b* [27]. Similarly, studies examining long-term memory in *Gadd45b*^*-/-*^ mutant mice have produced conflicting results. Sultan et al. (2012) found that knockout mice displayed enhanced coordination and balance on the accelerating rotarod, increased freezing following hippocampus-dependent contextual fear conditioning, and improved performance during the Morris water maze memory probe, together suggesting that *Gadd45b* may be negatively correlated with motor performance, fear learning, and spatial memory, respectively [27]. In contrast, studies by Leach et al. (2012) demonstrated decreased freezing in *Gadd45b*^*-/-*^ mice following contextual fear conditioning, proposing that *Gadd45b* plays an important role in hippocampus-dependent memory processes [40]. In both studies, *Gadd45b* manipulation had no effect on amygdala-dependent cued fear conditioning [27,40]. Whereas earlier studies involved transgenic knockout mice, the present study expands upon previous findings by implementing non-constitutive, brain region-specific knockdown of *Gadd45b* in the rat striatum using shRNA and CRISPR/Cas9 lentiviral vectors. The data presented here support a pro-memory role of *Gadd45b* in the striatum, with increases in *Gadd45b* mRNA following acute cocaine reward or exposure to a cocaine-paired environment (**Figure 1f**). Additionally, transgenic *Gadd45b*^*-/-*^ mice and adult rats with NAc-specific CRISPR/Cas9 mediated *Gadd45b* knockdown both exhibit drug-related memory deficits following cocaine-paired place conditioning (**Figure 2**). Furthermore, we demonstrate that *Gadd45b* knockdown results in differential upregulation of genes implicated in neurodegenerative diseases, such as Alzheimer’s disease (**Figure 4d**).

DNA methylation is a potent epigenetic regulatory modification that is critical for the function and information storage capacity of neuronal systems. In the brain, activity-dependent changes in DNA methylation are central regulators of synaptic plasticity and memory formation, and have been implicated in a broad range of neuropsychiatric disease states, including drug addiction. Previous studies have established that *Gadd45b* is required for activity-induced DNA demethylation in the hippocampus, specifically at CpG sites within regulatory regions of *Bdnf* and fibroblast growth factor-1 (*Fgf-1*), genes known to be important for adult neurogenesis in the dentate gyrus [21]. While no differences were observed in basal methylation levels within these gene regulatory regions in *Gadd45b* knockout mice, activity-induced DNA demethylation at these sites was nearly eliminated. In human post-mortem cortical tissue, decreased GADD45B binding to the *BDNF* promoter was associated with higher levels of 5mC and 5hmC [48]. In the mouse NAc, downregulation of *Gadd45b* alters DNA methylation in a phenotype-, gene-, and locus-specific way, potentially underlying susceptibility or resilience to stress [22]. Despite evidence suggesting that *Gadd45b* is important in activity-dependent DNA demethylation, it is still unclear how the GADD45B protein is involved in this process. GADD45B is an 18-kDa protein that belongs to the ribosomal protein L7Ae/L30e/S12e/Gadd45 superfamily and is known to interact with nuclear hormone receptors and bind nucleic acids [46,55]. GADD45B contains two LXXLL motifs, domains which are commonly found in other transcriptional coactivators and appear to be required for GADD45B to act as a transcriptional regulator [55]. Here, we examined DNA methylation profiles using RRBS following *Gadd45b* knockdown in striatal neurons. We demonstrate that baseline DNA methylation landscapes are preserved following *Gadd45b* knockdown (**Figure S2a-e**), but activity-induced changes in CpG methylation following DRD1 activation were absent in neurons lacking *Gadd45b* (**Figure S2f**). These results are consistent with prior studies, and suggest that *Gadd45b* is required for Drd1-dependent alterations in DNA methylation. Furthermore, while the above studies have implicated *Gadd45b* in activity-dependent DNA demethylation, we find that *Gadd45b* knockdown blocks both CpG hypermethylation (**Figure S2g-h**) and hypomethylation (**Figure S2i-j**) following treatment with SKF-38393. Finally, acute cocaine administration *in vivo* did not induce other genes known to be involved in DNA methylation processes in the NAc after 1 or 24 hours (**Figure 1b**). While our results are largely consistent with the known function of *Gadd45b* in stimulus-regulated DNA methylation changes, future experiments will be required to better characterize the molecular interactions that link *Gadd45b* to this process.

Using high-throughput electrophysiological approaches to record the activity of more than a thousand neurons *in vitro*, we observed that DRD1 receptor stimulation produced a significant increase in action potential frequency of striatal neurons (**Figure 5j-l**). This finding is consistent with recent discoveries that DA produces a rapid but sustained increase in excitability of D1 medium spiny neurons [56], as well as with prior results demonstrating a key role for DA in synaptic plasticity in this neuronal population [57]. Surprisingly, despite resulting in large-scale changes in gene expression, knockdown of *Gadd45b* did not significantly alter physiological changes in response to the DRD1 agonist SKF-38393. Instead, we observed that *Gadd45b* shRNA produced a relatively specific decrease in the duration of action potential burst events, without altering other electrophysiological parameters such as burst frequency, action potential frequency, or the number of spontaneously active neurons (**Figure 5e-i**). *In vivo*, burst firing events in medium spiny neurons are regulated by feedforward inhibition from local fast-spiking interneurons, and these events are especially critical for striatum-dependent learning and plasticity [58]. In freely moving rats, burst firing of medium spiny neurons in the NAc is temporally associated with important behavioral and reward-related events, including presentation of reward-paired cues [59,60], delivery/consumption of natural and drug rewards [60–62], and operant responses for rewarding stimuli [63–65]. Importantly, these phasic increases in neuronal activity encode key aspects of reward value and cost [66–68], and are thought to contribute to the key role of the NAc in associative learning and addiction [69,70]. Consistent with this role, we demonstrate that complete knockout or NAc-specific knockdown of *Gadd45b* also results in impaired cocaine-place learning. Future studies will be required to identify how *Gadd45b* contributes to specific learning-related activity of NAc neurons in the context of drug reinforcement.

In the current study, we describe a role for striatal *Gadd45b* in DA-dependent transcriptional regulation and cocaine reward, suggesting that *Gadd45b* is involved in the development and maintenance of addiction. These findings build upon previous work establishing a role for *Gadd45b* in neuropsychiatric disease. In patients with major psychosis, *GADD45B* mRNA expression was increased in post-mortem tissue from the inferior parietal lobule and the prefrontal cortex, corresponding with higher levels of GADD45B protein measured in cortical layers II, III, and V [48]. Despite the overall increase in protein, GADD45B binding is decreased at the *BDNF* promoter in patients with psychosis, corresponding to increased 5mC and 5hmC levels. In a rodent model of depression, *Gadd45b* expression was upregulated in the NAc of stress-susceptible mice following a chronic social defeat stress (CSDS) behavioral paradigm [22]. These increases in *Gadd45b* were not observed in stress-resilient mice, and downregulation of striatal *Gadd45b* rescued the social avoidance phenotype that distinguishes susceptible animals in CSDS. *Gadd45b* has also been implicated in neurodevelopmental diseases - *Gadd45b* expression is decreased in post-mortem cortical tissue from patients diagnosed with Autism Spectrum Disorder (ASD) [71]. Together, these results highlight that *Gadd45b* is dysregulated in the pathophysiology of other neuropsychiatric conditions. Although the underlying mechanisms of its involvement may differ by cell type and brain region, a thorough understanding of *Gadd45b*-dependent regulation of transcriptional and epigenetic dynamics will be an important next step for revealing potential therapeutic avenues targeting this pathway.

## Acknowledgements

This work was supported by NIH grants DP1-DA039650, R00-DA034681, and R01-MH114990 (J.J.D.), F32-DA041778 (F.A.S.), and T32-GM008361, and T32-GM008111 (M.E.Z.). L.I. is supported by the Civitan International Research Center at UAB. Additional assistance to J.J.D. was provided by the UAB Pittman Scholars Program. We thank Allison Bauman for generating our primary neuronal cell cultures. We thank Katherine Savell and Jasmin Revanna for their assistance with lentiviral production. We thank the Genomics Core Lab at the UAB Heflin Center for Genomic Sciences for assistance with RNA-seq and RRBS. We thank all current and former Day Lab members for assistance and support.

## Author contributions

J.J.D. performed RT-qPCR analysis of IEGs and DNA methylation machinery. A.C.B. completed cocaine locomotor sensitization assays and RT-qPCR analysis of *Gadd45b* expression after exposure to cocaine. F.A.S. cloned, designed, and validated CRISPR/Cas9 and shRNA constructs for *in vivo* and *in vitro* work, with assistance from G.G. J.J.D. and F.A.S. conceived of and performed CCP assays with transgenic mice. J.J.D., F.A.S., and M.E.Z. conceived of and performed CCP assays with CRISPR/Cas9 in rats, with assistance from G.G. M.E.Z. and F.A.S. completed all stereotaxic surgeries for *in vivo* testing, with assistance from G.G. and N.A.S. F.A.S. performed and analyzed *in vitro* DA, DRD1 agonist, and DRD2 agonist experiments with assistance from G.G. M.E.Z. performed and analyzed *in vitro* DRD1 antagonist, Forskolin, MEK inhibitor, and CREB inhibitor experiments. J.J.D., F.A.S., and L.I. performed statistical and graphical analysis of bulk RNA-seq datasets. M.E.Z. performed all electrophysiological assays and analyzed single-unit data, with assistance from J.J.D. and N.A.S. All projects were supervised by J.J.D. M.E.Z. and J.J.D. wrote the main text of the manuscript. All authors have approved the final version of the manuscript.

---

**Figure S1.**
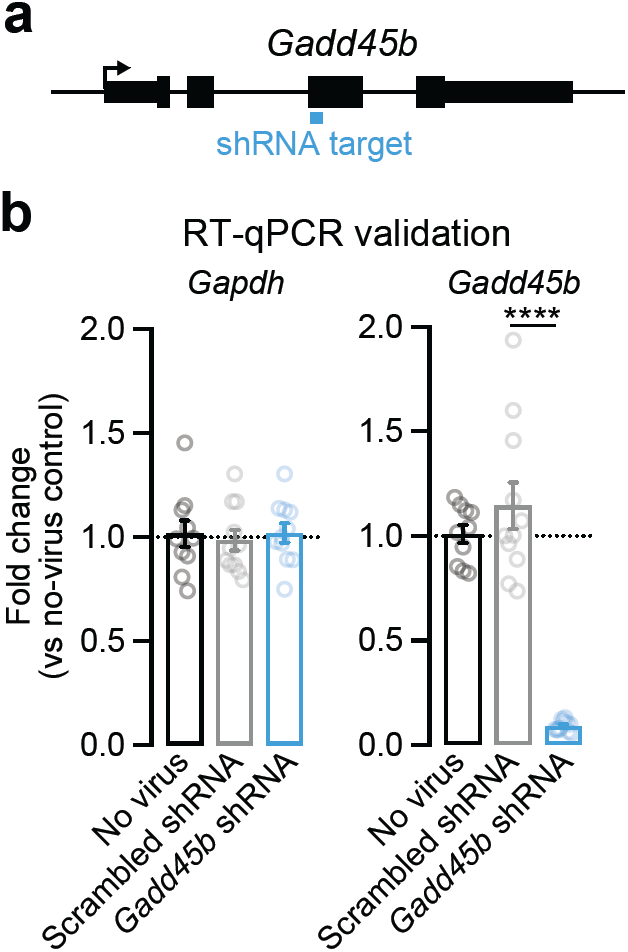
Efficient gene knockdown using RNA interference in primary striatal neuron cultures. **a**, A short hairpin RNA (shRNA) was engineered to target exon 3 of *Gadd45b* and packaged in a lentivirus backbone to drive gene knockdown. **b**, RT-qPCR validation revealed no effect on the housekeeping gene *Gapdh*, but a significant knockdown in *Gadd45b* mRNA compared to no-virus and scrambled shRNA controls (one-way ANOVA, *F*(2,29) = 66.82, *R*^*2*^ *=* 0.82, *p* < 0.0001; Tukey’s *post hoc* tests, \*\*\*\**p* < 0.0001 for indicated comparisons).

**Figure S2.**
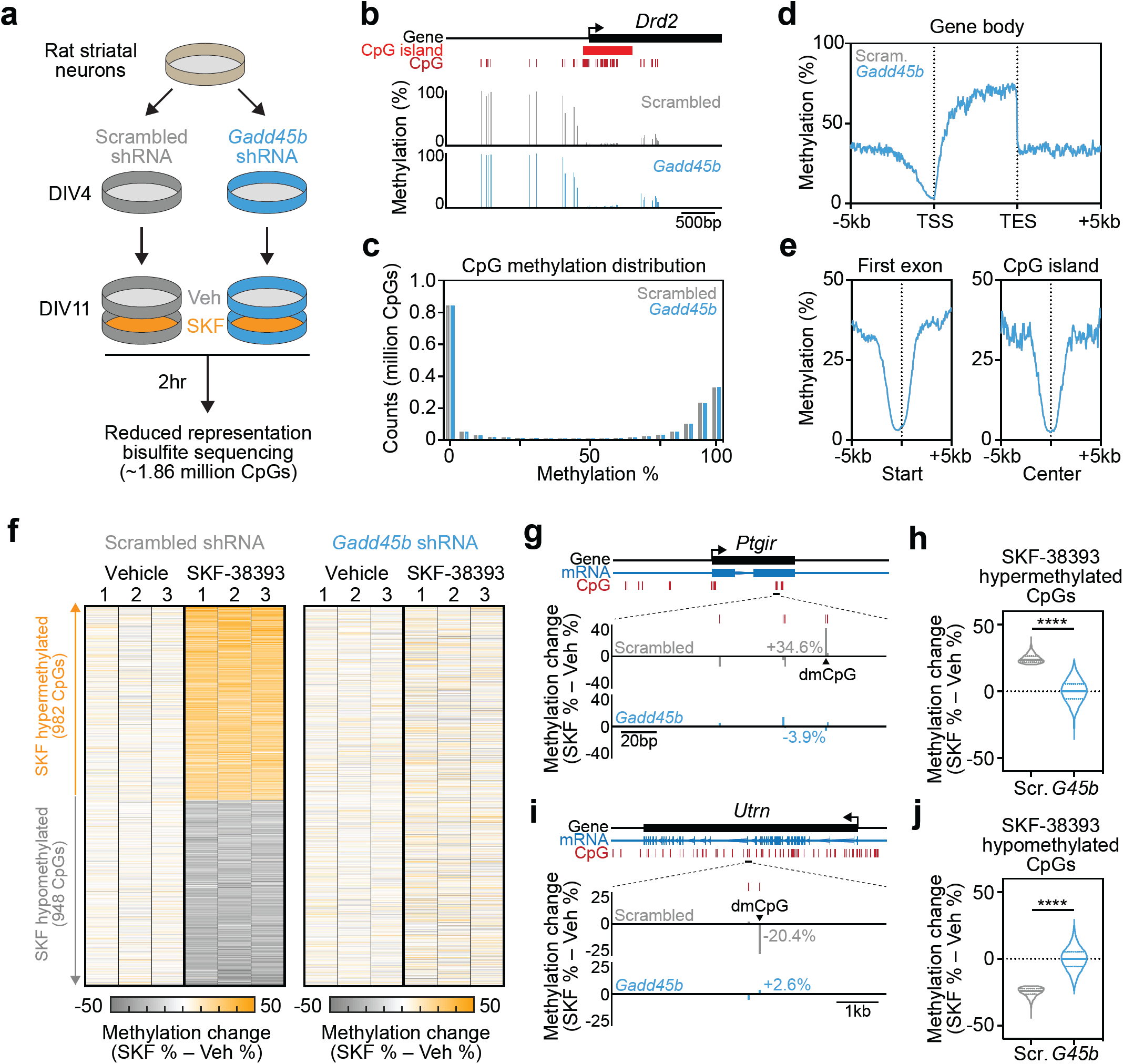
*Gadd45b* knockdown prevents dopamine-driven DNA methylation changes. **a**, Illustration of experimental design. Rat striatal cultures were transduced with *Gadd45b* shRNA (or scrambled control) at DIV4. Cultured neurons (*n* = 3/group) were treated with the DRD1 receptor agonist SKF-38393 (1μM) for 2hr at DIV11 prior to DNA extraction and genome-wide DNA methylation profiling with reduced representation bisulfite sequencing (RRBS). **b**, RRBS tracks from representative gene locus (*Drd2*) and promoter-spanning CpG island. Only CpGs with > 120 reads are shown. **c**, Genome-wide distribution of CpG methylation values from ∼1.86 million CpGs reveals bimodal distribution of CpG methylation in both scrambled and *Gadd45b* shRNA groups. **d**-**e**, DNA methylation profiles across gene bodies, first exon, and CpG islands reveals similar CpG methylation landscapes in scrambled and *Gadd45b* shRNA groups. **f**, Heatmaps showing CpG methylation change (SKF-38393 % - Veh %) for all 1930 CpGs modulated by SKF-38393 in the scrambled shRNA condition (termed differentially methylated CpGs (dmCpGs; defined as *p* < 0.01, > 20% change)). **g**, Representative SKF-38393 hypermethylated dmCpG at the *Ptgir* gene locus. **h**, *Gadd45b* shRNA prevents SKF-38393-induced increases in CpG methylation (Wilcoxon matched pairs signed-rank test, *W* = 449776, *p* < 0.0001). **i**, Representative SKF-38393 hypomethylated dmCpG at the *Utrn* gene locus. **j**, *Gadd45b* shRNA prevents decreases in CpG methylation following DRD1 agonist treatment (Wilcoxon matched pairs signed-rank test, *W* = −481671, *p* < 0.0001).

